# Is there more room to improve? The lifespan trajectory of procedural learning and its relationship to the between- and within-group differences in average response times

**DOI:** 10.1101/593582

**Authors:** Dora Juhasz, Dezso Nemeth, Karolina Janacsek

## Abstract

Characterizing the developmental trajectories of cognitive functions such as learning, memory and decision making across the lifespan faces fundamental challenges. Cognitive functions typically encompass several processes that can be differentially affected by age. Methodological issues also arise when comparisons are made across age groups that differ in basic performance measures, such as in average response times (RTs). Here we focus on procedural learning – a fundamental cognitive function that underlies the acquisition of cognitive, social, and motor skills – and demonstrate how disentangling subprocesses of learning and controlling for differences in average RTs can reveal different developmental trajectories across the human lifespan. Two hundred-seventy participants aged between 7 and 85 years performed a probabilistic sequence learning task that enabled us to separately measure two processes of procedural learning, namely general skill learning and statistical learning. Using raw RT measures, in *between*-group comparisons, we found a U-shaped trajectory with children and older adults exhibiting greater general skill learning compared to adolescents and younger adults. However, when we controlled for differences in average RTs (either by using ratio scores or focusing on a subsample of participants with similar average speed), only children (but not older adults) demonstrated superior general skill learning consistently across analyses. Testing the relationship between average RTs and general skill learning *within* age groups shed light on further age-related differences, suggesting that general skill learning measures are more affected by average speed in some age groups. Consistent with previous studies of learning probabilistic regularities, statistical learning showed a gradual decline across the lifespan, and learning performance seemed to be independent of average speed, regardless of the age group. Overall, our results suggest that children are superior learners in various aspects of procedural learning, including both general skill and statistical learning. Our study also highlights the importance to test, and control for, the effect of average speed on other RT measures of cognitive functions, which can fundamentally affect the interpretation of group differences in developmental, aging and clinical psychology and neuroscience studies.

## Introduction

Procedural learning is a fundamental cognitive function that facilitates efficient processing of and automatic responses to complex environmental stimuli, supporting efficient adaptation to the changing environment. Procedural learning underlies the acquisition of new cognitive, social, and motor skills (Lieberman, 2000; Nemeth et al., 2011; Romano Bergstrom, Howard, & Howard, 2012; Ullman, 2016); it is therefore a critical function across the human lifespan. It is a widely held view that procedural learning is most effective in childhood; nevertheless, acquiring new skills such as learning languages or learning to use new devices and applications are also possible throughout adulthood. These lifelong learning abilities are increasingly sought out in the workplace as they contribute to economic competitiveness. Additionally, it has been shown that at least in some cases learning new skills can serve as a shield against age-related cognitive decline (Bialystok, Craik, Binns, Ossher, & Freedman, 2014; Park et al., 2014; Perani et al., 2017). Despite the ubiquitous nature of procedural learning throughout the human lifespan, how learning is affected by age is not yet fully understood.

Previous research on age-related changes in procedural learning reported mixed findings. Pioneering studies found comparable performance in children and young adults (Karatekin, Marcus, & White, 2007; Meulemans, Van der Linden, & Perruchet, 1998; Saffran, Aslin, & Newport, 1996), while more recent evidence suggests age-related differences in learning. Some of these more recent studies showed that young adults outperform children (Hodel, Markant, Van Den Heuvel, Cirilli-Raether, & Thomas, 2014; Lukács & Kemény, 2015; Thomas et al., 2004), while others found better learning performance in childhood than in adults (Fischer, Wilhelm, & Born, 2007; Janacsek, Fiser, & Nemeth, 2012; Nemeth, Janacsek, & Fiser, 2013). Aging studies also yielded heterogeneous results, with older adults exhibiting weaker (Bennett, Howard, & Howard, 2007; J. H. Howard, Jr. & Howard, 1997; Schuck et al., 2013), comparable (Gaillard, Destrebecqz, Michiels, & Cleeremans, 2009; Lukács & Kemény, 2015; Spencer, Gouw, & Ivry, 2007), or in some cases even better (Brown, Robertson, & Press, 2009) learning performance than younger adults. The theoretical frameworks of age-related changes also reflect this heterogeneity. 1) The developmental invariance model claims age independence in procedural learning (Reber, 1993). 2) The inverted U-shape model argues that a peak in learning performance should occur in young adulthood (Thomas et al., 2004). 3) The ‘children better’ model claims that children should exhibit superior learning performance compared to adolescents and adults (Janacsek et al., 2012; Newport, 1990). Thus, in summary, both empirical findings and theoretical frameworks present a puzzle of age-related differences in procedural learning. A possible solution to this puzzle may lie in taking into account the multifaceted nature of learning. Here, we aim to disentangle different processes underlying procedural learning that can contribute to a better understanding of age-related differences in skill learning.

Procedural learning is a multifaceted cognitive function that encompasses multiple processes. A critical process within procedural learning is how our brain extracts and learns the structures of the environment, including repeated sequences and occurrence statistics of perceptual stimuli (typically termed as sequence-specific and statistical learning, respectively) (Gaillard et al., 2009; Kóbor et al., 2018; Simor et al., 2019; Turk-Browne, Scholl, Chun, & Johnson, 2009). Here we refer to learning these structures as *task-specific learning* since the to-be-learned structures may be specific to a given environment, and more specifically, to a given task. In contrast to task-specific learning, some more general processes also contribute to procedural learning, including faster processing of and responding to the perceptual stimuli, and faster matching of the corresponding responses to those stimuli (i.e., improved visuomotor coordination) as the task progresses (Hallgato, Győri-Dani, Pekár, Janacsek, & Nemeth, 2013; Kóbor, Janacsek, Takács, & Nemeth, 2017). We refer to these processes as *task-general learning* or general skill learning as these processes may be relatively independent of the specific stimulus structure. Disentangling task-specific and task-general effects has posed a challenge to numerous procedural learning tasks (such as in finger sequence tapping and in variants of the Serial Reaction Time (SRT) task; Pan & Rickard, 2015; Robertson, 2007), and a failure to separately assess these processes may at least partially explain the contradictory findings of age-related differences in procedural learning.

To demonstrate how task-specific and task-general learning can contribute to procedural learning, take an example of learning to drive a car. In this case, task-general or general skill learning includes efficient perceptual processing of our environment (e.g., road signs, and other cars on the road), a fast motor system to plan and execute movements, and efficient coordination between the perceptual and motor system. In contrast, task-specific or sequence-specific learning, involves the serial ordering (sequencing) of several actions during driving; for instance, when waiting at the red light that turns green (processing of a visual stimulus) we press the clutch pedal with the left foot (motor response), then shift gear with the right hand, and finally press the gas with the right foot. Clearly, the efficient visuomotor processing and coordination is as critical for successful driving as is knowing and performing the planned actions in the appropriate serial order. Since these lower level processes show clear age-related differences (e.g., children are generally slower than adolescents and young adults, Janacsek et al., 2012; Thomas & Nelson, 2001), it is reasonable to assume that age-related differences also emerge in task-general learning, which relies on these lower level processes. Consequently, if various tasks depend on task-specific vs. task-general learning processes to a different degree and their contributions to performance cannot be disentangled in these tasks, this may explain a significant portion of the mixed findings of age-related differences in procedural learning.

How can age-related differences in these lower level (perceptual-motor) processes affect task-general and task-specific learning? It has been suggested that slower responses generally provide ‘more room to improve’ during practice (e.g., Bhakuni & Mutha, 2015; Jiang, Capistrano, Esler, & Swallow, 2013). According to this argument, participants with initially slower response times should exhibit a greater speed-up (i.e., task-general learning) during practice compared to others with initially faster response times. Since children and older adults typically show slower responses than younger adults, this should result in greater task-general learning in the former two age groups. The predictions for how age-related differences in lower level processes should affect task-specific learning are less clear. It is possible that participants with generally slower response times exhibit greater task-specific learning, although this possibility seems more plausible if task-general and task-specific learning processes cannot be properly teased apart. Testing these possibilities can have important consequences for developmental and aging studies as they can lead to fundamentally different interpretations by disentangling genuine age-related differences in procedural learning from purely methodological (measurement) issues. Importantly, however, these possibilities have not yet been directly and systematically tested, particularly from a lifespan perspective.

In the present study, we use a procedural learning task – the Alternating Serial Reaction Time (ASRT) task – that enables us to distinguish between task-specific and task-general learning (Csabi, Varszegi-Schulz, Janacsek, Malecek, & Nemeth, 2014; Janacsek & Nemeth, 2012) in order to separately probe their developmental trajectories and their relationships with average speed across the human lifespan. In this four-choice perceptual-motor reaction time task, visual stimuli appear on the screen following an alternating sequential pattern that is repeatedly presented throughout the task (J. H. Howard, Jr. & Howard, 1997; Nemeth et al., 2010). Participants are required to respond to these stimuli by pressing the corresponding response keys on a keyboard as fast and as accurate as they can. The repeating alternating sequence results in some stimulus combinations (so-called triplets) being more frequent than others, and during practice participants learn to differentiate between these more frequent and less frequent stimulus combinations (often referred to as triplet or statistical learning; Janacsek, Borbély-Ipkovich, Nemeth, & Gonda, 2018; Kóbor et al., 2017; Unoka et al., 2017). At the same time, independent of task-specific learning, participants become generally faster as the task progresses (often referred to as general skill learning). This general speed-up during practice may be attributed to the processes described above, including faster processing of the visual stimuli, and faster matching of the corresponding response keys to those stimuli as the task progresses (Hallgato et al., 2013; Kóbor et al., 2017). Thus, in the ASRT task, triplet or statistical learning captures *task-specific learning*, and the general speed-up during task captures *task-general learning*.

Procedural learning across the human lifespan has been previously probed with the ASRT task in one study, focusing on the age-related differences in task-specific learning (Janacsek et al., 2012). This study showed a gradual decline across the lifespan in triplet learning with best learning performance in children. Additionally, a U-shaped developmental trajectory of average speed (i.e., RTs averaged for the entire learning session) was also reported: consistent with previous observations, children and older adults exhibited generally slower responses than adolescents and young adults. This study, however, did not directly test task-general learning and the relationship between average speed and learning. Here, we aim to address these gaps in a new sample of participants (*N* = 270) aged between 7 and 85 years.

The aims of the present study are thus twofold. First, we aimed to examine age-related differences in general skill learning across the human lifespan. Second, we also aimed to test the argument of whether generally slower RTs are associated with greater general skill learning in different developmental stages across the lifespan. We used several approaches to achieve these aims focusing both on between-group and within-group differences. Namely, age-related differences in general skill learning were tested using both raw RT learning scores as well as ratio scores that can potentially control for differences in average RTs across age groups. Additionally, an alternative approach is also presented whereby age-related differences in general skill learning are compared in a subsample of participants whose average RTs are similar across age groups. While these approaches explore between-group differences in general skill learning and its potential relationship with average speed, direct evidence for such relationship can be better obtained from within-group comparisons. To this end, we performed correlation analyses between average RTs and general skill learning indices (both raw RT and ratio scores) separately for each age group. Finally, we also report age-related differences in triplet learning and test for a relationship between average RTs and task-specific learning. Beyond allowing for a comparison of how average RTs may affect task-general vs. task-specific learning, the presentation of triplet learning results provides a direct replication test of Janacsek et al.’s (2012) study, promoting reproducible research.

## Materials and methods

### Participants

Two hundred-seventy participants aged between 7 and 85 years took part in the experiment. They were assigned to nine age groups (*n* = 30 in each group). Fourteen participants were excluded based on their slower response times or lower accuracy during the whole experiment (3 *SD*s outliers) compared to their respective age group. The final sample consisted of 256 participants. Mean and standard deviation for age and education, and gender ratio for all age groups is presented in Table 1. None of the participants suffered from any developmental, psychiatric or neurological disorders. All participants gave signed informed consent (parental consent was obtained for children), and they received no financial compensation for their participation. The study was approved by the local university ethics committee and was conducted in accordance with the Declaration of Helsinki.

**Table 1.** Demographic data (means, standard deviations, and proportions) for all age groups.

| **Group** | **Age** | **Gender** | **Education** |
| --- | --- | --- | --- |
| 7-8-year-old (n=26) | 7.92 (0.27) | 13 M / 13 F | 2.00 (0.00) |
| 9-10-year-old (n=28) | 9.79 (0.42) | 13 M / 15 F | 4.00 (0.27) |
| 11-13-year-old (n=30) | 12.10 (0.61) | 13 M / 17 F | 6.00 (0.00) |
| 14-15-year-old (n=30) | 14.55 (0.57) | 13 M / 17 F | 8.10 (0.40) |
| 16-17-year-old (n=30) | 16.56 (0.54) | 13 M / 17 F | 10.12 (0.45) |
| 18-29-year-old (n=30) | 21.64 (2.93) | 12 M / 18 F | 14.52 (2.05) |
| 30-44-year-old (n=30) | 36.67 (3.81) | 12 M / 18 F | 14.73 (2.93) |
| 45-60-year-old (n=26) | 51.65 (4.46) | 6 M / 20 F | 13.56 (4.62) |
| 61-85-year-old (n=26) | 66.04 (5.84) | 5 M / 21 F | 12.92 (3.92) |

### Task and procedure

We used the ASRT task (J. H. Howard, Jr. & Howard, 1997; Nemeth et al., 2010) where a stimulus (a dog’s head) appeared in one of the four empty circles arranged horizontally on a computer screen. Participants were instructed to respond to the stimulus events by pressing the corresponding response keys (Z, C, B or M on a QWERTY keyboard) as fast and accurately as they could. The ASRT task consisted of 20 blocks with 85 key presses in each block. The first five responses of each stimulus block served for practice only (and were excluded from the analyses), and then the eight-element alternating sequence (e.g., 2R1R3R4R, where numbers refer to the four locations on the screen, and ‘R’ refers to a randomly selected location out of the four possible ones) was repeated ten times within a block. The stimulus remained on the screen until participants pressed the correct response key, and the next stimulus appeared 120 ms after the correct response. Between blocks, participants received feedback on the screen about their overall reaction time (RT) and accuracy. The computer program generated a different repeating ASRT sequence of the four locations for each participant using a permutation rule such that each of the six unique permutations of the four repeating events occurred with equal probability (for more details see e.g., Kóbor et al., 2017; Szegedi-Hallgató et al., 2017).

In this study we focused on the following measures derived from the ASRT task: 1) *average RTs*, defined as the average speed across the entire task; 2) *general skill learning*, defined as the RT change from the beginning to the end of the task; and 3) *triplet learning*, defined as faster responses for more frequent stimuli compared to the less frequent ones (see e.g., Kóbor et al., 2017).

### Statistical analysis

Data preprocessing, calculation of the ASRT measures of interest and statistical analyses followed the procedures outlined in previous ASRT studies (D. V. Howard et al., 2004; J. H. Howard, Jr. & Howard, 1997; Kóbor et al., 2017; Nemeth et al., 2010; Song, Howard, & Howard, 2007). Briefly, the 20 blocks of the ASRT task were organized into four segments (called *epochs*), each consisting of five blocks (i.e., Blocks 1-5 corresponds to Epoch 1, Blocks 6-10 corresponds to Epoch 2, etc.). We calculated the median RTs of correct responses separately for high- and low-frequency triplets, for each epoch and for each participant.

First, to test age-related differences in general skill learning (and triplet learning) across age groups, the raw RT values were submitted to a mixed-design ANOVA with EPOCH (1 to 4) and TRIPLET TYPE (high-vs. low-frequency) as the within-subject factor, and GROUP (9 age groups) as the between-subject factor. In this ANOVA, the main effect of EPOCH can reveal general skill learning, and the EPOCH x GROUP interaction can reveal differences in general skill learning across age groups.

Second, to test age-related differences in general skill learning while controlling for group differences in average RTs, we calculated general skill learning ratio scores. Namely, for each participant, the average RT of Epoch 4 was subtracted from the average RT of Epoch 1 (which provides the raw RT difference score of general skill learning) and then divided by the average RT of that participant in the entire task. An advantage of such a ratio score is that it can be interpreted as a percentage change in RTs during practice relative to one’s average speed. This ratio score was submitted to a Univariate ANOVA with GROUP (9 age groups) as the between-subject factor.

For all ANOVAs, Greenhouse-Geisser epsilon (ε) correction was used when necessary. Original *df* values and corrected *p* values (if applicable) are reported together with partial eta-squared (*η*_*p*_ ^2^) as the measure of effect size. Post-hoc analysis was conducted by Fisher’s LSD pairwise comparisons.

Third, to gain a better understanding of between- and within-group heterogeneity, we also present individual data for average RTs and general skill learning indices (raw RT difference, ratio score), with quadratic model fitting. Additionally, the relationship between average RTs and learning measures was tested by Pearson correlation analyses. These analyses can reveal whether the argument of ‘slower RTs provide more room to improve’ is supported *within* age groups and whether the relationship between average RTs and general skill learning changes *across* age groups. Moreover, these analyses can also reveal whether (and in which age groups) the ratio scores is adequate for controlling for average RT differences.

Fourth, we also explored an alternative, complementary approach to test the potential age-related differences in general skill learning while controlling for the average RTs. Namely, we selected a subsample of participants that exhibited similar average RTs, irrespective of their age groups. Thus, we asked whether participants in different age groups exhibit comparable degree of general skill learning if their average RTs are similar. To put it differently: if a 9-year-old child is as fast as a 16-year-old adolescent, do they exhibit a similar degree of general skill learning? Due to the fact that young adults are, on average, faster than children and older adults, this selection criterion necessarily leads to the inclusion of faster participants in children and older adult groups and slower participants in the young adult groups relative to their respective age groups. Nevertheless, it still may provide valuable information for the relationship between age and general skill learning while controlling for the average RTs.

Fifth, we also report age-related differences in triplet learning based on the main ANOVA described above (focusing on the main effect of TRIPLET and TRIPLET x GROUP interactions; (Janacsek et al., 2012; Nemeth, Janacsek, Király, et al., 2013) and test for a relationship between average RTs and triplet learning (using Pearson correlation analyses). As described in the Introduction, beyond providing a comparison of how average RTs may affect task-general vs. task-specific learning, we find the presentation of triplet learning results important as it can provide a direct replication test of Janacsek et al.’s (2012) study, promoting reproducible research.

## Results

### Are there age-related differences in average RT and general skill learning?

The mixed-design ANOVA revealed a significant main effect of GROUP (*F*(1, 247) = 28.858, *η*_*p*_ ^2^ = 0.483, *p* < .001) as the average RTs differed significantly across age groups (Figure 1A). The LSD post hoc test revealed gradually faster RTs between 7 and 15 years of age (*p*s < .023), similarly fast RTs between 15 and 29 years of age (*p*s > .600), and then gradually slower RTs between 30 and 85 years of age (*p*s < .060). These results confirm a U-shaped developmental trajectory in average RTs across the lifespan (Janacsek et al., 2012).

**Figure 1.**
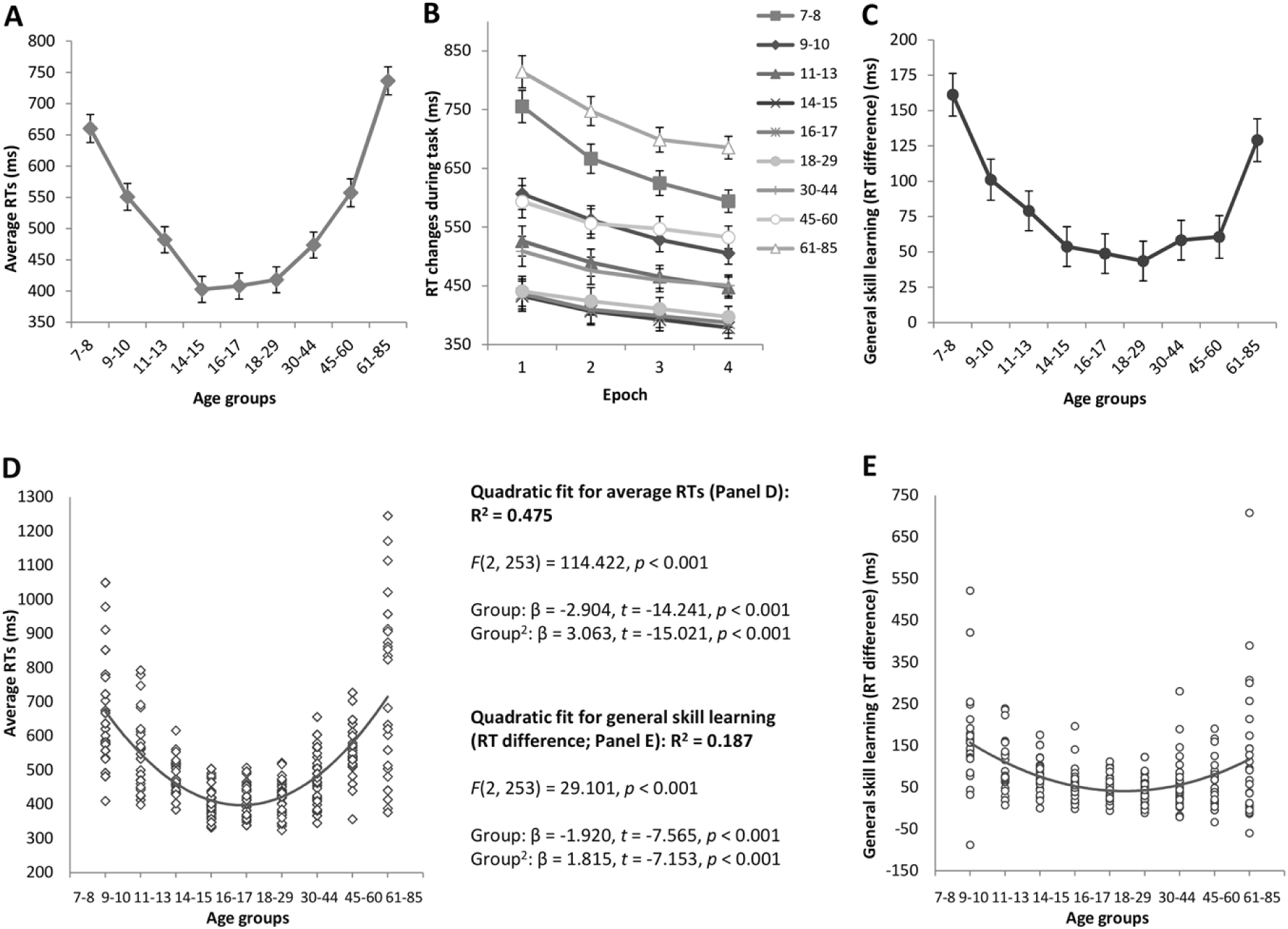
Average RTs and general skill learning across the lifespan. Average RTs refer to the RTs of all correct responses averaged over the entire task (A). General skill learning refers to the RT changes occurring during the time course of the task (B), which can be quantified as the RT difference between Epoch 1 and Epoch 4 (C). Group averages are presented in panel A-C, and individual data separately for each age group are presented in panel D-E. Error bars indicate standard error of mean (SEM).

Regarding general skill learning, the ANOVA revealed a significant main effect of EPOCH (*F*(3, 741) = 191.197, *η*_*p*_ ^2^ = 0.436, *p* < .001), with significantly faster RTs as the task progressed. More importantly, the EPOCH × GROUP interaction was also significant (*F*(24, 741) = 5.484, *η*_*p*_ ^2^ = 0.151, *p* < .001), suggesting significantly different general skill learning across age groups (Figure 1B). The LSD post-hoc test comparing the RT differences between Epoch 1 and 4 across age groups revealed that the 7-8-year old age group exhibited the greatest general skill improvement (Figure 1C), significantly differing from all other age groups (*p*s < 0.004) except for the 61-85-year old group (*p* = .134). The 9-10- and 11-13-year-old groups showed a smaller improvement, with no difference between the two groups (*p* =.277). From adolescence to late adulthood, the degree of general skill learning further decreased compared to the younger age groups, with no group differences between 14 and 60 years of age (*p*s > .409). The 61-85-year-old group’s general skill learning differed significantly from that of the groups between 11 and 60 years of age (*p*s < .016).

To gain a better understanding of between- and within-group differences, individual data for each participant in each age group are presented in Figure 1D and 1E. In line with the ANOVA results, a quadratic fit for the average RTs explains a large proportion (47.5%) of variability across age groups (Figure 1D). Interestingly, the quadratic fit for the general skill learning (Figure 1E) is substantially weaker compared to the average RT fit, explaining only 18.7% of variability across age groups. After excluding the three slowest participants from the 7-8 and 61-85-year-old groups, the fit is only slightly better, 20.9%, still well below of that for the overall RTs. (Note that excluding these participants affected the ANOVA results of general skill learning reported above only in that the 7-8-year-olds showed significantly greater learning compared to all age groups, including the 61-85-year-olds as well, *p*s <.023).

Both the developmental trajectories of group averages (Figure 1AC) as well as the quadratic fits for average RTs and general skill learning (Figure 1DE) seem to support the argument that if someone is slower, then there is more room to improve during practice. Those age groups that exhibit slower average RTs seem to show greater general skill learning (i.e., more speed-up from Epoch 1 to Epoch 4). Nevertheless, the substantially weaker quadratic fit for general skill learning compared to the average RTs suggests that (between- or within-group) differences in average RTs alone may not be sufficient to explain differences in general skill learning. In the next steps, first we focus on between-group differences, and then we will test the within-group associations between average RTs and general skill learning.

### Are there age-related differences in general skill learning when average RT differences are controlled for?

The Univariate ANOVA on the general skill ratio score yielded a significant main effect of GROUP (*F*(8, 247) = 4.602, *η*_*p*_ ^2^ = 0.130, *p* < .001), suggesting differences in general skill learning across age groups (Figure 2A). Based on the LSD post hoc test, the 7-8-year-old group showed the highest general skill ratio score (23.6% improvement relative to their average RTs) that was significantly different from that of all other age groups (*p*s < .043). The 9-10- and 11-13-year-old groups exhibited a smaller improvement (17.9% and 16.2%, respectively), with no difference between the two groups (*p* = .532). The degree of general skill learning exhibited a further decrease after that age, and remained comparable between 14 and 85 years of age (*p*s > 0.126; except for the 18-29 vs. 61-85-year-old groups: *p* = .057).

**Figure 2.**
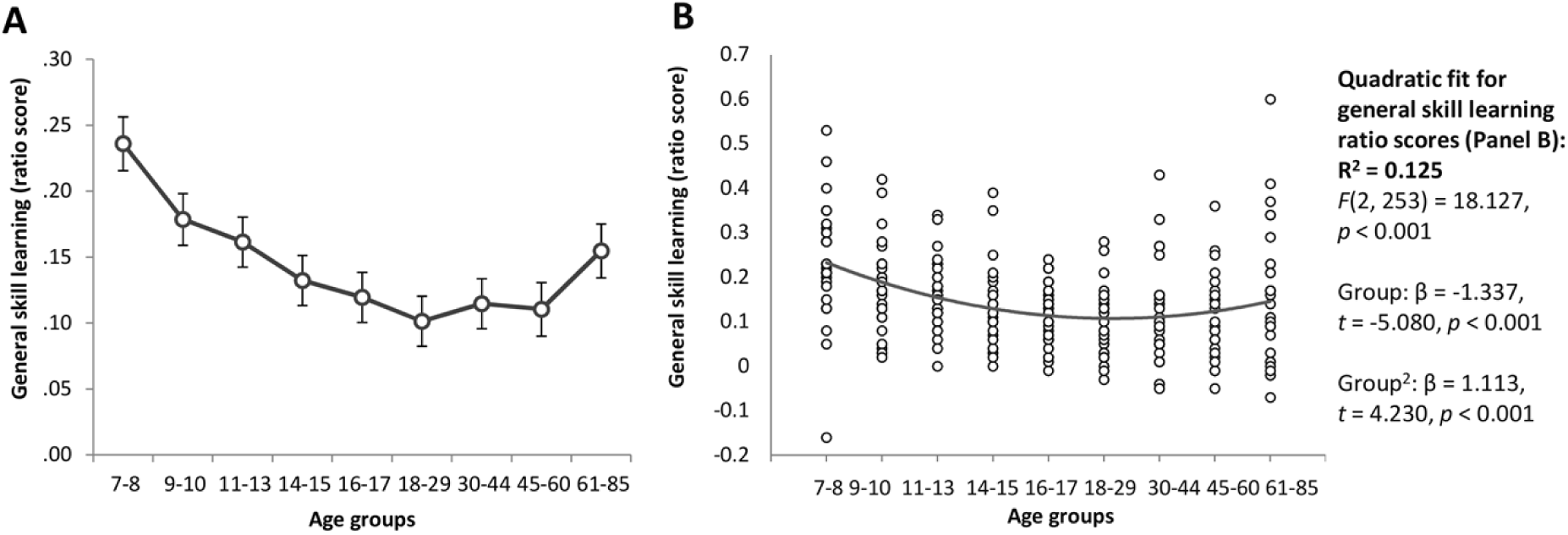
General skill learning ratio scores across the lifespan. The ratio scores for group averages (A) and individual data (B) are presented. The ratio score can be interpreted as a percentage change in performance (e.g., the 7-8-old age group exhibited an approximately 23% speed-up from Epoch 1 to Epoch 4 relative to their average speed during task). Error bars indicate standard error of mean (SEM).

The individual data of the ratio scores are presented in Figure 2B. Although the quadratic fit for the general skill ratio scores was weaker than that for the average RTs and for the raw RT difference of general skill learning, it still explained 12.5% of variability across age groups. This is in line with the ANOVA results of significant group differences, existing mainly between children (particularly 7-8-year-olds) and the other age groups.

Comparing the raw RT difference and the ratio score as measures of general skill learning, a considerable difference can be observed in their developmental trajectories: While the 61-85-year-old group seems to have greater general skill learning than the groups between 11 and 60 years of age based on the raw RT difference between Epoch 1 and Epoch 4 (Figure 1C), this greater improvement largely disappears when the ratio score is used (there is only a trend level difference compared to the 18-29-year-old group). This result suggests that in the 61-85-year-old group the observed RT changes during the task may be more affected by the participants’ average speed compared to the other age groups, and thus may reflect other processes as well, over and beyond those related to general skill learning.

A similar approach to control for differences in average RTs is to use the average RT of Epoch 1 instead of the average RT of the entire task when calculating the ratio scores. This approach has been previously used in the study of procedural learning to control for RT differences across groups (e.g., Bhakuni & Mutha, 2015; Nitsche et al., 2003). In this case, performance at the beginning of the task (i.e., in Epoch 1) is set to the same level for all participants (the value of 1) and performance changes during the remaining of the task (i.e., in Epochs 2 to 4 in this case) are relative changes compared to the performance in Epoch 1. This score can also be interpreted as a percentage change similar to the one used above. We re-ran our analysis with this score and obtained almost identical results as the ones reported above (with small differences in numerical values only).

### Are larger average RTs associated with greater general skill learning within age groups?

In the next step, we tested the relationship between average RTs and the degree of general skill learning for each age group separately. The Pearson correlation analyses revealed a moderate positive relationship between average RTs and the raw RT difference of general skill learning between 7 and 13 years of age, between 18 and 44 years of age, as well as in the 61-85-year-old group (Table 2). This relationship was eliminated in the groups between 7 and 13 years of age when the ratio score of general skill learning was used, suggesting that dividing the RT change during the task by the average RT can adequately control for the effect of average speed on the speed-up during the task, and thus may provide a less biased measure of general skill learning for group comparisons. However, the ratio score proved less effective to control for this relationship in adulthood as the average RTs still remained positively correlated with the general skill ratio score in the 30-44-year-old group (and on trend level in the 18-29- and 61-85-year-old groups). It is also important to note that there was no relationship between average RTs and general skill learning (either the raw or the ratio score) in the 14-15 and 16-17-year-old groups, suggesting that the relationship itself may change in different developmental stages. Similarly, there was no relationship between average RTs and general skill learning in the 45-60-year-old group either.

**Table 2.** Relationship between average RTs and indices of general skill (raw RT difference and ratio scores) and triplet learning for each age group.

| Group | General skill learning (raw RT difference) | General skill learning (ratio score, relative to average RT) | Triplet learning |
| --- | --- | --- | --- |
| 7-8-year-old | $r = .582, p = .002$ | $r = .260, p = .200$ | $r = .233, p = .253$ |
| 9-10-year-old | $r = .556, p = .002$ | $r = .258, p = .184$ | $r = .237, p = .225$ |
| 11-13-year-old | $r = .476, p = .008$ | $r = .306, p = .101$ | $r = .158, p = .405$ |
| 14-15-year-old | $r = .302, p = .105$ | $r = .127, p = .502$ | $r = -.043, p = .820$ |
| 16-17-year-old | $r = .238, p = .205$ | $r = .036, p = .852$ | $r = .020, p = .917$ |
| 18-29-year-old | $r = .416, p = .022$ | $r = .319, p = .085$ | $r = .189, p = .317$ |
| 30-44-year-old | $r = .624, p < .001$ | $r = .500, p = .005$ | $r = .293, p = .116$ |
| 45-60-year-old | $r = -.007, p = .973$ | $r = -.120, p = .558$ | $r = .136, p = .508$ |
| 61-85-year-old | $r = .546, p = .004$ | $r = .381, p = .055$ | $r = .057, p = .781$ |

*Note.* Pearson correlation coefficients are reported. Significant correlations ( $p < .05$ ) are highlighted with a darker gray background, and trend level correlations ( $p < .1$ ) are highlighted with a lighter gray background.

Overall, based on the correlation analysis, there appears to be a positive relationship between average RTs and raw RT changes during task in most age groups, and the ratio score of general skill learning can help decrease or eliminate this relationship.

### Do participants in different age groups exhibit comparable degree of general skill learning if their average RTs are similar?

To answer this question, based on the individual data of each age group (cf. Figure 1D), we selected a subsample of participants whose average RTs were between 400 and 550 ms. The RT range seemed to provide an appropriate balance between maximizing the sample size in each age group and minimizing the potential average RT differences in group averages. Table 3 shows to what percentiles of the original sample the selected range of 400-550 ms corresponds.

**Table 3.**
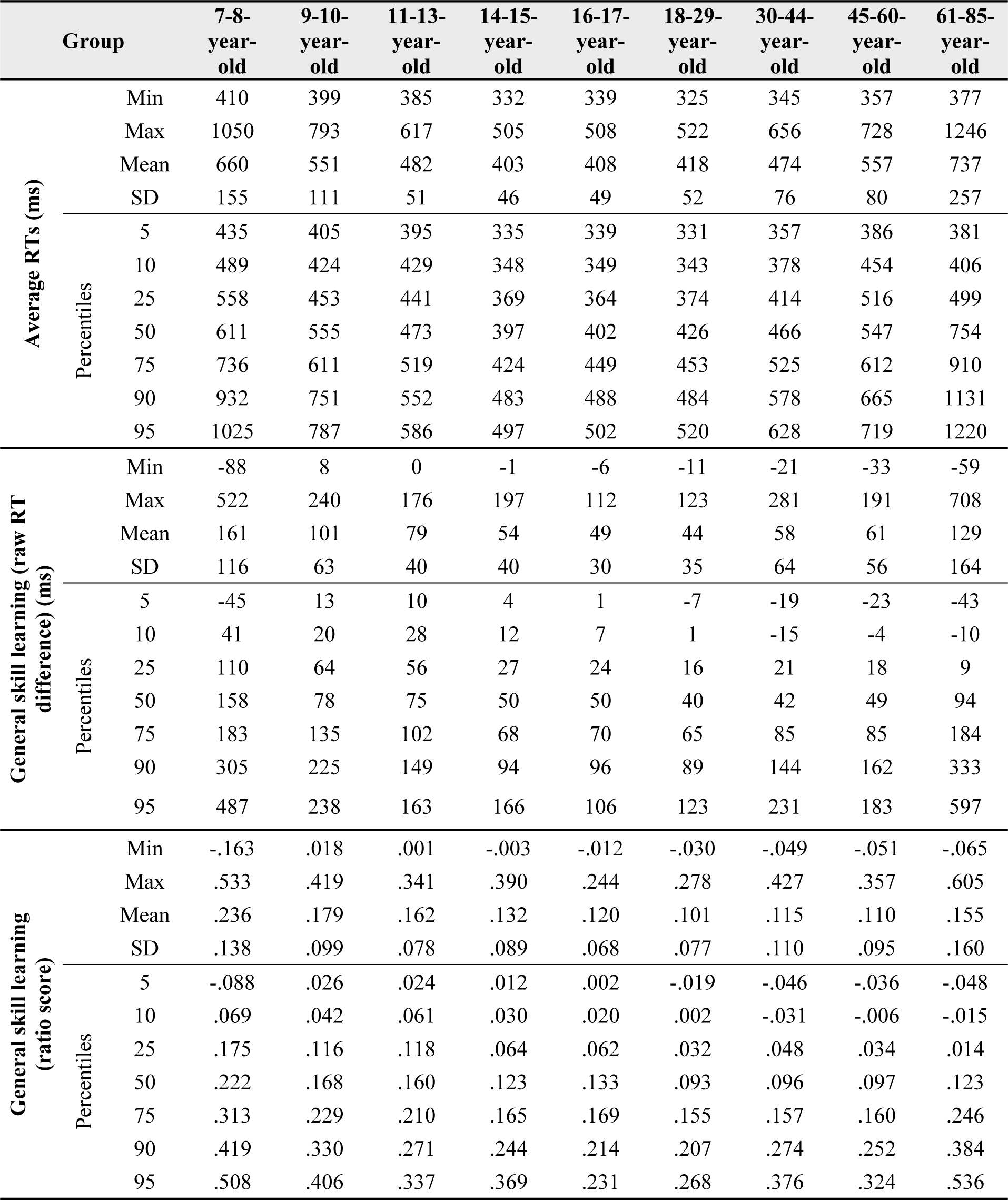
Minimum and maximum values, mean, standard deviation (SD) and percentiles of average RTs and general skill learning for each age group.

Since the number of participants by groups is relatively low and unequal, standard statistical analyses may be less reliable here. Therefore, in Figure 3 we present the same group averages for this subsample as the ones presented in Figure 1A-C and Figure 2A for the whole sample in order to enable qualitative comparison between the pattern of results. Figure 3A suggests that, with this approach, group differences in average RTs can be at least partly eliminated. Importantly, some age differences in general skill learning still remained (Figure 3B-D). To support the qualitative interpretation of these results, we compared the ratio score across age groups, focusing primarily on the 7-8- and 61-85-year-olds. This analysis revealed greater general skill ratio scores for the 7-8-year-olds compared to the 61-85-year-olds (*p* =.035). Additionally, the 61-85-year-old group’s general skill ratio score did not differ significantly from those between 14 and 60 years of age (*p*s > .125). Although this statistical analysis should be treated carefully because of the low and unbalanced sample sizes across groups, it is important to highlight that the results presented here (both qualitatively and quantitatively) support the interpretation of age-related differences in general skill learning obtained in the whole sample. Specifically, children (particularly 7-8-year-olds) seem to show better general skill learning compared to later ages, and this advantage cannot be explained by their overall slower response speed, as it persists even after controlling for the average RTs. In contrast, older adults may not show better general skill learning, and the observed greater RT changes during task may be due to other factors, including overall slower response speed, as their advantage diminishes when average RTs are controlled for.

**Figure 3.**
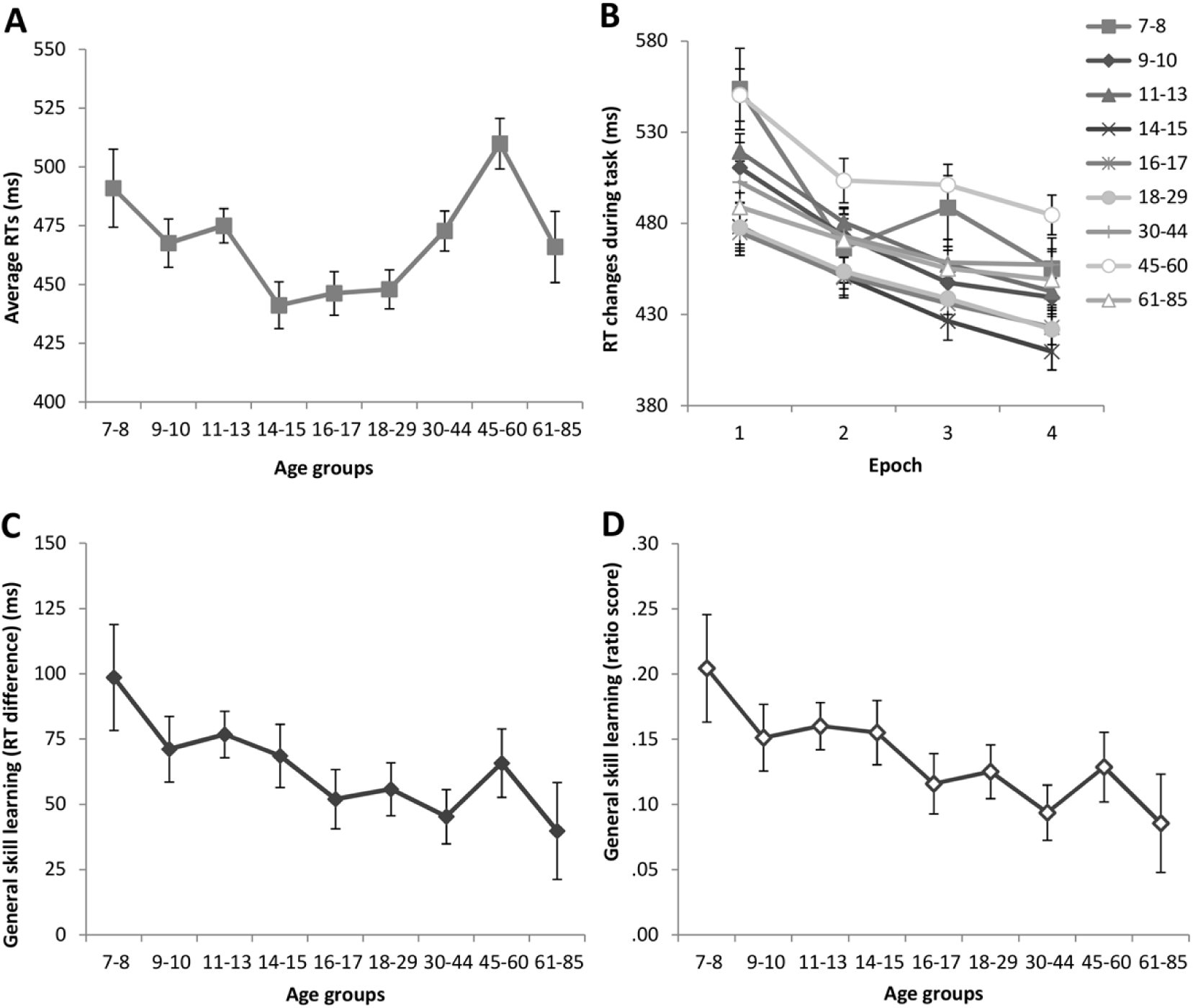
Average RTs (A) and general skill learning (B-D) across the lifespan for a subsample of the participants to control for age-related average RT differences. Only those participants are included in this subsample whose average RTs are between 400 and 550 ms. For details on the measures presented here see the legend of Figure 1. Error bars indicate standard error of mean (SEM).

### Are there age-related differences in triplet learning? Additionally, **is there a relationship between triplet learning scores and average RTs?**

Although it is not of the main interest of the current paper, we briefly report the results of triplet learning as well (Figure 4). The main ANOVA (described in the *Statistical analysis* section) revealed a significant a main effect of TRIPLET (*F*(1, 247) = 228.365, *η*_*p*_ ^2^ = 0.480, *p* < .001), with significantly faster RTs for more frequent triplets compared to the less frequent ones. The TRIPLET × GROUP interaction was also significant (*F*(8, 247) = 2.598, *η*_*p*_ ^2^ = 0.078, *p* = .010). The LSD post hoc test revealed a pattern similar to the one reported in the Janacsek et al. (2012) study, with similar triplet learning performance in the 7-13 age range (*p*s >.490) that was significantly higher than the learning scores in the 14-85 age range (*p* < .001).

**Figure 4.**
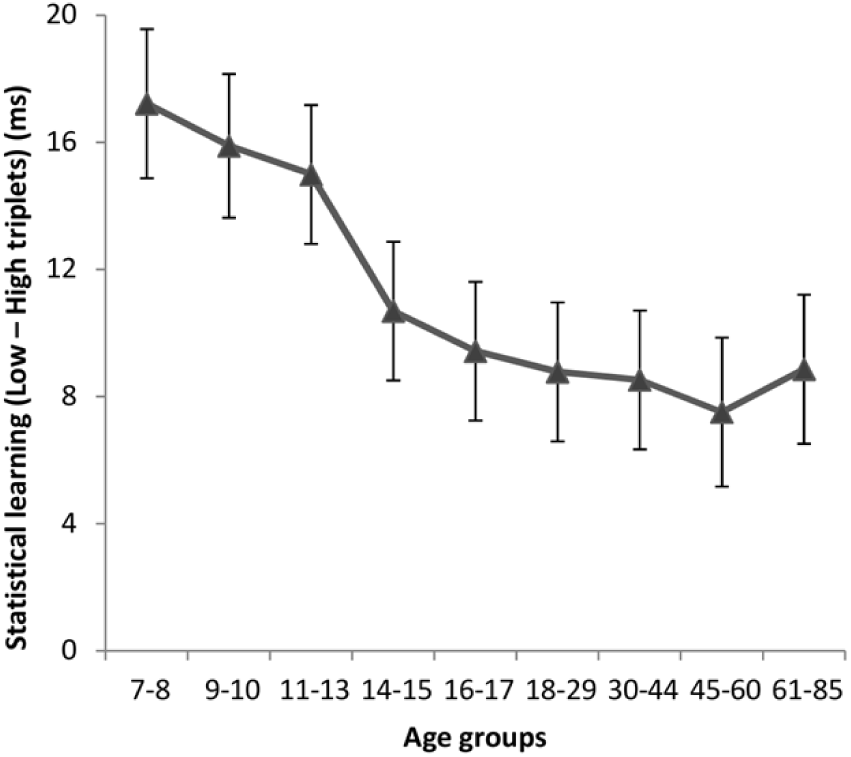
Statistical learning across age groups. Triplet learning score was quantified as an RT difference for low- and high-frequency triplets, averaged across the entire task. Larger values represent better learning performance. Error bars indicate standard error of mean (SEM).

Additionally, we tested the relationship between average RTs and triplet learning scores in each age group separately, and found no significant correlation between these measures in either age group (Table 2). Thus, it seems that while general skill learning may be correlated with average RTs in some developmental stages, supporting the claim of ‘slower RTs, thus more room to improve’, triplet learning scores appear to be unrelated to average RTs.

## Discussion

Our study aimed to examine age-related differences in general skill learning across the human lifespan and test the argument of whether generally slower response times are associated with greater general skill learning in different developmental stages. We employed the ASRT task, which probes procedural learning, and enables us to disentangle task-general (i.e., general skill) learning from task-specific learning of probabilistic regularities (i.e., statistical learning). A large sample of participants aged between 7 and 85 years were tested on this task. We found a U-shaped developmental trajectory of general skill learning, assessed by raw RT changes during the task, with children and older adults exhibiting greater learning than adolescents and young adults. This developmental trajectory was paralleled with a U-shaped lifespan trajectory of average RTs, lending support to the ‘more room to improve’ argument from a between-group perspective. Nevertheless, our more detailed analyses of both between-group and within-group differences suggest a more complex relationship between average speed and general skill learning across the human lifespan. Importantly, the superior general skill learning of children (particularly that of the 7-8-year-olds) was consistently demonstrated across different analysis approaches, while the older adults’ general skill learning decreased to the level of young adults when differences in average speed were controlled for. Finally, task-specific triplet learning showed a gradual decline across the lifespan, and learning performance seemed to be independent of average speed, regardless of the age group. Overall, our results suggest that children are superior learners in various aspects of procedural learning, including both task-specific and task-general processes of learning.

We used three different approaches to test the lifespan trajectory of general skill learning. Using raw RT measures, we observed a U-shaped trajectory with children and older adults exhibiting greater general skill learning compared to adolescents and younger adults. In contrast, when we used RT ratio scores (which is a common approach to control for differences in average speed across groups; see e.g., Nitsche et al., 2003; Urry, Burns, & Baetu, 2018), the developmental trajectory was no longer U-shaped. Our results showed an advantage for children compared to adults, while the older adults (the 61-85-year-old group) no longer exhibited greater general skill learning compared to the younger adults, suggesting that the greater speed-up observed in raw RT measures may due to different factors in children vs. older adults, even if both age groups show slower average speed compared to young adults. Additionally, as a third approach, we tested age-related differences in a subsample of participants whose average RTs were similar across age groups. Even though the selection of such subsample may have induced some bias (as faster participants of the children and older adult groups and slower participants of the adolescent and young adult groups were included in this analysis), it still may help provide a deeper insight into age differences in general skill learning while controlling for average RT differences. This analysis further confirmed the ratio score results of the whole sample: the 7-8-year-old group exhibited the greatest general skill learning (approximately 20%, comparable to the ratio score of the whole sample of this age group), and then gradually smaller improvements were observed from childhood to adulthood. The 61-85-year-old group did not exhibit greater general skill learning than any other adult group. Thus, overall, children (especially the 7-8-year old age group) exhibited superior general skill learning consistently across analyses. This finding suggests a heightened ability to acquire new skilled behaviors in this developmental stage that cannot be explained by the generally slower responses in childhood.

It is a typically observed pattern in developmental, aging and clinical neuroscience and psychology studies that groups with different average speed (e.g., children/older adults vs. young adults, or patient vs. control groups) are compared on different RT measures (Bennett, Romano, Howard, & Howard, 2008; Gordon & Stark, 2007; Janacsek et al., 2012; Negash et al., 2007; Takács et al., 2017). In these group comparisons the difference in average speed poses a great challenge to disentangle its effect from other RT measures, such as those related to learning (e.g., speed-up as learning progresses), in order to unravel the more fine-grained group differences in cognitive functions. To better understand the extent of this challenge, here we tested the argument of whether generally slower response times are associated with greater general skill learning (i.e., slower average RTs provide ‘more room to improve’) in different developmental stages. The between-group comparisons showed a similar U-shaped lifespan trajectory both for average speed and general skill learning measured by raw RTs, which can easily be viewed as support for the ‘more room to improve’ argument. However, our more detailed analyses of both between-group and within-group differences suggest a more complex relationship between average speed and general skill learning across the human lifespan. In between-group comparisons, as discussed above, children showed superior general skill learning performance compared to other age groups even when average speed across groups was controlled for, either by using ratio scores or in the analysis of a subsample with similar average speed. These results suggest that differences in average speed alone cannot, at least fully, explain the observed differences in general skill learning across the lifespan.

Moreover, when we tested the relationship between average speed and general skill learning *within* age groups, we found correlations in age groups 7 to 13, 18 to 44, and 61 to 85 years but not in adolescents (aged 14 to 17 years) or in middle aged participants (aged 45 to 60 years), suggesting that the relationship between average speed and general skill learning (measured by raw RT differences) is not universal. Importantly, the use of ratio scores for general skill learning appeared to efficiently control for the differences in average speed in children as the correlation between average speed and general skill learning disappeared in the 7-13-year-old age groups when such ratio scores were used. In contrast, ratio scores seemed less efficient in controlling for average speed differences in adulthood (particularly in the 30-44-year-old group). These findings have important implications for developmental, aging and clinical studies comparing groups with different average speed: they highlight the importance of testing the relationship between average speed and other RT measures of interest, and the need for finding measures that can efficiently control for differences in average speed while not inducing other biases in group comparisons.

In the current study we also found age-related differences in task-specific triplet learning: the lifespan trajectory resembled a gradual decline across age groups with children exhibiting the best learning performance. This finding provides a direct replication of the Janacsek et al. (2012) study in a new, independent sample of participants, suggesting, that at least in learning probabilistic regularities, children show an advantage compared to other age groups. This result supports the ‘children better’ theoretical framework in contrast to other models that emphasize developmental invariance or peak learning performance in young adulthood (Fletcher, Maybery, & Bennett, 2000; Meulemans et al., 1998; Reber, 1993; Thomas & Nelson, 2001). Additionally, the current study enabled us to test, in the same sample of participants, whether average speed is similarly related to general skill learning vs. triplet learning in order to gain a better understanding of the relationship between various aspects of procedural learning. Interestingly, while average speed showed a positive relationship with general skill learning (see above), no such relationship was observed with triplet learning in any of the age groups. This may emerge from the fact that triplet learning scores are computed as difference scores between RTs for high-vs. low-frequency triplets (J. H. Howard, Jr. & Howard, 1997; Nemeth et al., 2010). Thus, if someone is, on average, slower than others, s/he will show slower responses to *both* triplet types throughout the task, but the RT difference to these triplets (i.e., how much faster they respond to high-vs. low-frequency triplets) seems to be independent of the average speed of participants. This result is in line with findings of Török et al. (2017) showing that such triplet learning measures are resistant to factors that affect general performance changes, such as fatigue effects (Török, Janacsek, Nagy, Orbán, & Nemeth, 2017). Thus, overall, the triplet learning measure appears to be a well-designed, reliable tool for testing the learning of probabilistic regularities (Stark-Inbar, Raza, Taylor, & Ivry, 2016; Török et al., 2017), and is well-suited for group comparisons, even if average speed differs across groups. In contrast, other RT measures (e.g., general skill learning measures) may be more sensitive to average speed differences across participants and groups.

Although we found better learning performance in children, it is important to note that children do not universally show an advantage compared to adults (Lukács & Kemény, 2015; Zwart, Vissers, Kessels, & Maes, 2017), and other factors should also be taken into account to gain a better understanding of procedural learning across the lifespan. Such factors may include the structure to be learned in the task (e.g., triplets/statistics, probabilistic or deterministic sequences) (Kóbor et al., 2018; Lukács & Kemény, 2015; Simor et al., 2019; Stark-Inbar et al., 2016), their presentation parameters (e.g., simultaneous or sequential) (Janacsek & Nemeth, 2012), stimulus timing, task length, fatigue effects (Pan & Rickard, 2015; Török et al., 2017), or the ratio of motor vs. perceptual components of learning (Deroost & Soetens, 2006; Hallgato et al., 2013; Nemeth, Hallgato, Janacsek, Sandor, & Londe, 2009). Future studies should systematically test these potential factors. Our study highlights the importance of using tasks that are able to assess different aspects of procedural learning, including the differentiation of performance changes that are related to general skill learning (i.e., task-general learning) vs. those related to learning the structure embedded in the task (i.e., task-specific learning). Moreover, the relationship between learning measures and average speed should also be tested and appropriately controlled for, when comparing groups with different average speed. Such research approach could significantly advance our understanding of differences in procedural learning across the lifespan.

Additionally, our study has important implications for a wide range of clinical populations, including neurodevelopmental (e.g., dyslexia, autism, ADHD, Tourette Syndrome) and neurodegenerative disorders (e.g., Mild Cognitive Impairment, Alzheimer’s disease) as well as psychiatric conditions (e.g., schizophrenia, depression, bipolar disorder). These clinical populations typically exhibit slower responses compared to the healthy/typically developing control groups (Bennett et al., 2008; Gordon & Stark, 2007; Janacsek et al., 2018; Negash et al., 2007; Scheu et al., 2013; Schmitter-Edgecombe & Rogers, 1997; Schwartz, Howard, Howard, & Hovaguimian, 2003; Takács et al., 2017). Understanding the relationship between average speed and aspects of procedural learning in different developmental stages can help formulate predictions for and test how various clinical conditions alter these processes, taking age-related differences into account as well. In the current study, we reported detailed parameters of average speed and general skill learning for each age group separately that can be used as reference in future clinical studies.

Here we focused on reaction time measures and their developmental trajectories across the lifespan. Although it is out of the scope of the current paper, it is important to note that accuracy measures can also be analyzed in at least some tasks of procedural learning. The patterns of overall accuracy and changes in accuracy during learning may exhibit different developmental trajectories and different within-group relationships compared to the reaction time measures. For example, children tend to have lower, whereas older adults tend to have higher overall accuracy than young adults, thus, average accuracy and average speed seems to show different lifespan trajectories (D. V. Howard et al., 2004; Janacsek et al., 2012; Meulemans et al., 1998). It is reasonable to assume that the mechanisms related to average accuracy as well as to the changes in accuracy during learning are, at least partly, different from the mechanisms related to average speed and its changes (speed-up) during learning. Accuracy measures may be more closely related to attention and action selection functions, such as selectively attending to the target stimuli and selecting the appropriate responses to those stimuli. In contrast, reaction time measures usually include correct responses only, and thus, the attentional and action selection processes may affect these measures to a smaller extent. Instead, the average speed and the speed-up due to practice typically observed in procedural learning tasks may be more closely related to achieving greater automaticity that is a hallmark of skilled behaviors (Ashby & Crossley, 2012; Logan, 1988). Future studies should directly test these possible mechanisms and their differential contributions to accuracy and reaction time measures.

In summary, children (especially the 7-8-year-olds) exhibited superior general skill learning consistently across analyses, suggesting a heightened ability to acquire new skilled behaviors in this developmental stage, which extends the ‘children better’ theoretical framework of procedural learning (Janacsek et al., 2012) to include both task-general and task-specific processes. Our study highlights the importance of disentangling these processes of procedural learning as they may be differentially affected by age or clinical conditions (see e.g., Bennett et al., 2008; Csabi et al., 2014; Meulemans et al., 1998) and may be differentially related to average speed. Here we presented two approaches to control for average speed differences across groups: using ratio scores and testing a subsample of participants with similar average speed. Overall, the argument that slower average speed provides ‘more room to improve’ seems to be not universally true: some age groups across the lifespan and some measures of learning (task-general but not task-specific learning) seem to be more affected by average speed. Thus, our findings highlight the importance to test, and control for, the effect of average speed on other RT measures of cognitive functions, which can fundamentally affect the interpretation of group differences in developmental, aging and clinical psychology and neuroscience studies.

## Acknowledgments

This research was supported by the National Brain Research Program (project 2017-1.2.1-NKP-2017-00002); Hungarian Scientific Research Fund (OTKA PD 124148, to K. J.; NKFIH-OTKA K 128016, to D. N.); Janos Bolyai Research Fellowship of the Hungarian Academy of Sciences (to K. J.).

